# Direct Ink Writing of Graphene Oxide Reinforced 13-93B1 Bioactive Glass Scaffolds for Bone Tissue Engineering Applications

**DOI:** 10.1101/2024.03.14.584172

**Authors:** Kartikeya Dixit, Ashok Vishwakarma, Hitendra Kumar, Keekyoung Kim, Niraj Sinha

## Abstract

Graphene-reinforced bioactive glass scaffolds have gained significant attention in the field of bone tissue engineering due to their unique combination of mechanical strength, bioactivity, and electrical conductivity. Additive manufacturing techniques, such as 3D printing, provide a versatile platform for fabricating these scaffolds with precise control over their architecture and composition. Consequently, in this work, we have fabricated graphene oxide (GO)-reinforced 13-93B1 bioactive glass scaffold using the direct ink writing technique. A Pluronic F-127- based ink was prepared for scaffold fabrication, and its rheological properties were assessed for shear thinning behaviour, structural support, and recovery. Further, the fabricated scaffolds were characterized using micro-computed tomography, scanning electron microscopy, and energy dispersive x-ray spectroscopy. Additionally, computational fluid dynamics simulations with Dulbecco’s modified eagle medium and blood were performed to evaluate the perfusion kinetics of the scaffolds. The inclusion of GO enhanced the compressive strength of the fabricated scaffolds by ∼225%. The morphological characterization based on micro-computed tomography showed that additively manufactured scaffolds have appropriate porosity, pore size, pore throat size, and interconnectivity. The live-dead assay results showed no cytotoxicity towards C2C12 mouse myoblast cells. Also, cell adhesion and cell viability results show better cell growth on the nanocomposite scaffolds. Overall, the fabricated scaffold is found suitable for bone tissue engineering applications.

## 1. Introduction

Segmental bone defects caused by trauma, tumor removal, and congenital disorders are prevalent clinical concerns. While micro-fractures in the bone can heal over time, segmental defects require external intervention, such as bone grafting. Bone autograft, bone allograft, and prosthetic implants are currently commonly utilized to repair significant bone lesions [1]. However, they have certain limitations. For instance, autologous bone grafts have hindrance such as limited availability and donor site morbidity. On the other hand, bone allografts and prosthetic implants have unpredictable long-term durability and expensive prices [2]. Under these conditions, porous synthetic scaffolds that replicate bone would be ideal bone substitutes. Still, they must sustain tissue ingrowth and integration with the native bone and surrounding soft tissues. Ideally, they must degrade at a rate consistent with new bone development.

The first silicate bioactive glass, 45S5, has been employed in most biomedical research and applications since it was discovered by Hench et al. in 1971 [3]. More recently, 13-93, a silicate bioactive glass composed of 53 SiO_2_, 6 Na_2_O, 12 K_2_O, 5 MgO, 20 CaO, and 4 P_2_O_5_, has gained attention for its potential in the medical field. The Food and Drug Administration of the United States has approved the *in vivo* usage of 45S5 and 13-93 glass. However, 45S5 and 13-93 bioactive glasses have certain limitations as scaffold materials for bone repair and regeneration. Sintering 45S5 particles into a porous 3D network with sufficient strength for healing bone lesions is challenging due to the glass’s inclination to crystallize before considerable viscous flow. In addition, the conversion rate of 45S5 and 13-93 glass to hydroxyapatite (HA) in the simulated body fluid is slow and partial [4,5].

It is well known that bioactive glasses encourage the growth of bone cells and form strong bonds with hard and soft tissue, making them ideal for filling bone deficiencies. Bioactive glasses, once implanted, transform into a calcium phosphate or HA material, forming a strong attachment with the native tissue. In addition to releasing ions that activate the expression of osteogenic markers, bioactive glasses have also been demonstrated to stimulate angiogenesis. Graphene is a viable substitute for reinforcing scaffolds. On the one hand, graphene makes it possible to change the structure of materials at the nanometre scale to make stronger and tougher bioceramics. Further, it has recently been observed that graphene can facilitate faster cell growth and division [6]. However, this is still debated and requires more detailed studies. Therefore, using graphene as a strengthening phase in additively manufactured scaffolds could improve their mechanical properties and biological performance at the same time. Previously, researchers have demonstrated that incorporating reduced graphene oxide (rGO) into additively manufactured 45S5 bioglass scaffolds enhances their compressive strength and toughness [7]. However, there are challenges to achieving a homogeneous dispersion of rGO in the bioglass matrix. Due to its hydrophobic nature, rGO can agglomerate together in the aqueous ink suspensions used in additive manufacturing, dropping the composites’ mechanical characteristics [8]. Therefore, to replace rGO as a reinforcement for additively manufactured bioactive glass scaffolds, graphene oxide (GO) could be employed due to higher dispersibility in water [9].

In addition to directly mediating cell adhesion, proliferation, and differentiation by promoting fibronectin extracellular protein adsorption and enriching various growth factors in body fluids, it has been widely demonstrated that GO can cause physical and chemical damage to bacteria through high surface tension and oxidative stress [10,11].

The optimal scaffold materials must effectively utilize osteoinduction, osteoconduction, and osseointegration in order to optimize bone regeneration. Consequently, the multicomponent scaffolds exhibit potential as viable options for bone tissue engineering. In this work, we prepared the composite of 13-93 B1 (18B_2_O_3_–36SiO_2_– 22CaO–2P_2_O_5_–6Na_2_O–8MgO–8K_2_O (mol%)) and GO. GO acted as a reinforced component. Various weight percentages of GO were dispersed in a bioactive glass matrix. The scaffolds were fabricated by using direct ink writing technique. Compressive strength of the fabricated scaffolds was evaluated. Cellular behaviour of the GO reinforced scaffolds was evaluated using *in vitro* cell culture studies.

## 2. Materials and Methods

### 2.1. Materials

The following chemicals were used for synthesis of 13-93B1 bioactive glass: tetraethyl orthosilicate (Si(OC_2_H_5_)_4_, purity >98%), triethyl phosphate ((C_2_H_5_O)_3_PO, purity >99.8%), calcium nitrate tetrahydrate (Ca(NO_3_)_2_.4H_2_O, purity >98%), boric acid (H_3_BO_3_, purity >99.5%), sodium nitrate (NaNO_3_, purity 99%), magnesium nitrate hexahydrate (Mg(NO_3_)_2_. 6H_2_O, purity 97%), potassium nitrate (KNO_3_, purity 99.5%) and nitric acid (HNO_3_, 69% assay). For graphene oxide synthesis, graphite powder, sulfuric acid (H_2_SO_4_), phosphoric acid (H_3_PO_4_), potassium permanganate (KMnO_4_) and hydrogen peroxide (H_2_O_2_) were used. The aforementioned chemicals were procured from Merck India.

### 2.2. Synthesis of bioactive glass

The sol-gel method starts with a 90-minute hydrolysis step wherein 0.55 mol of Si(OC_2_H_5_)_4_ was mixed with 1 M HNO_3_ and deionized water (please refer to Figure 1). Next, 0.03 mol of (C_2_H_5_O)_3_PO was added after hydrolysis, followed by sequential addition of 0.34 mol of Ca(NO_3_)_2_.4H_2_O, 0.09 mol of NaNO_3_, 0.12 mol of Mg(NO_3_)_2_. 6H_2_O, 0.12 mol of KNO_3_, and 0.28 mol of H_3_BO_3_. The time of reaction between each addition of the chemical was 30 minutes. Once the sol was prepared, it was placed into a glass bottle and maintained at 70 °C in a hot air oven for three days to facilitate gel formation. Next, the oven temperature was increased to 150 °C for two days to dry the gel. The bioactive glass was then stabilized at 600 °C to eliminate nitrates added during its manufacturing. The bioactive glass powders were ground using a ball mill for 48 hours using ethanol (C_2_H_2_OH, 99.9% purity) as the milling medium to generate fine particles (5–10 µm). The resulting powders were dried in a hot air oven to eliminate ethanol.

**Figure 1.**
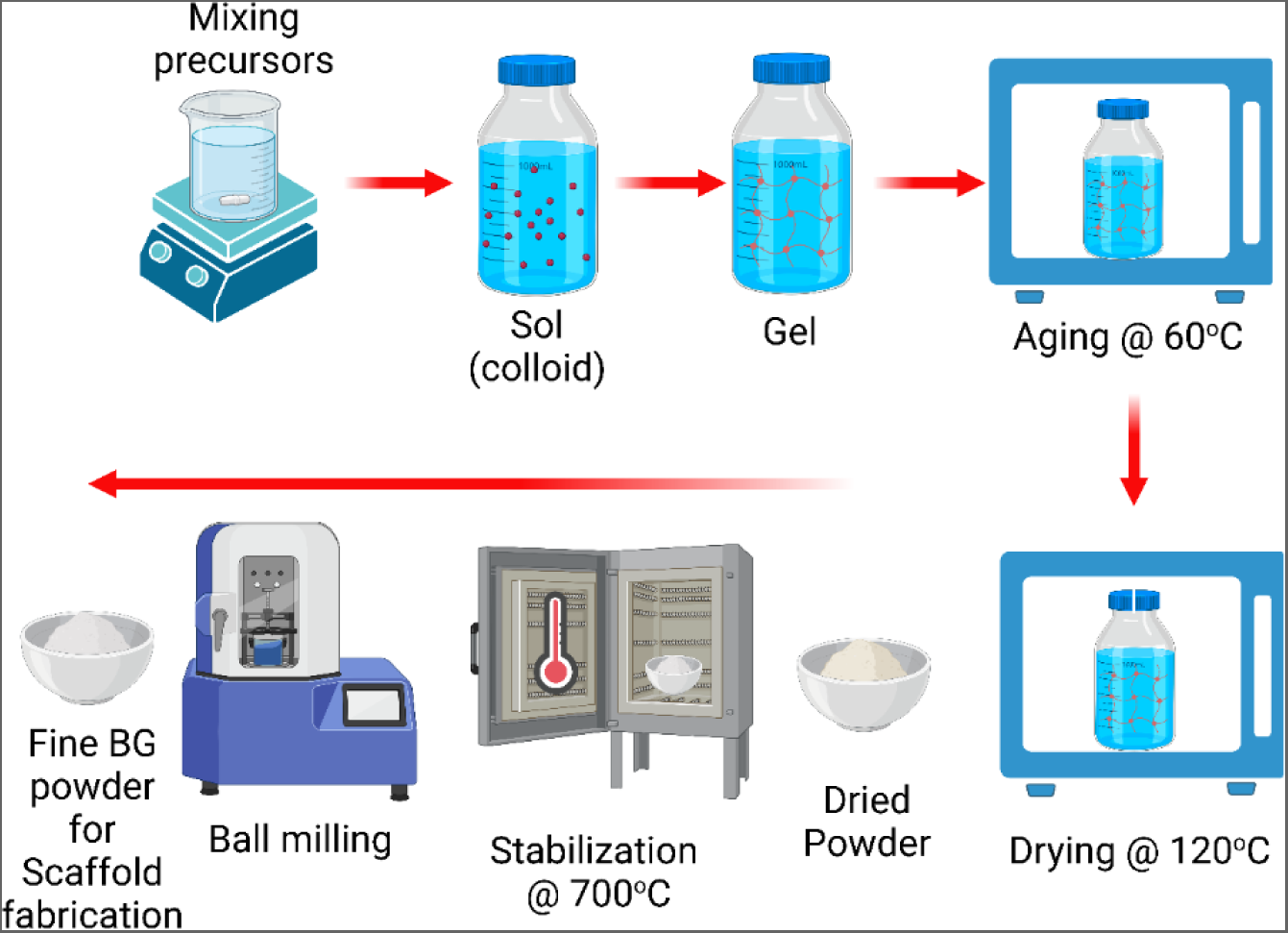
Schematic showing sol-gel synthesis method of 13-93B1 bioactive glass.

### 2.3. Synthesis of Graphene Oxide

The modified Hummer’s method was used to synthesize the graphene oxide (GO) in this study (please refer to Figure 2) [12]. Initially, 1g graphite powder was thoroughly mixed with 81 mL and 9 mL concentrated sulfuric acid (H_2_SO_4_) and phosphoric acid (H_3_PO_4_), respectively using a magnetic stirrer at 500 rpm for 3 hours. Afterwards, 4 g potassium permanganate (KMnO_4_) was gradually added to the solution. The whole solution was mixed until the color of solution changed to dark green indicating the oxidation of graphite flakes. The excessive KMnO_4_ was neutralized with dropwise addition of 30% H_2_O_2_ into the solution. Next, the solution was filtered using filter paper. The collected precipitate was washed with 20 mL HCl and 1 L deionized (DI) water to remove the impurities. Further, the collected GO was repeatedly washed with DI water until the pH increased to 7. The obtained GO was dispersed in DI water using ultrasonication to ensure homogeneous dispersion.

**Figure 2.**
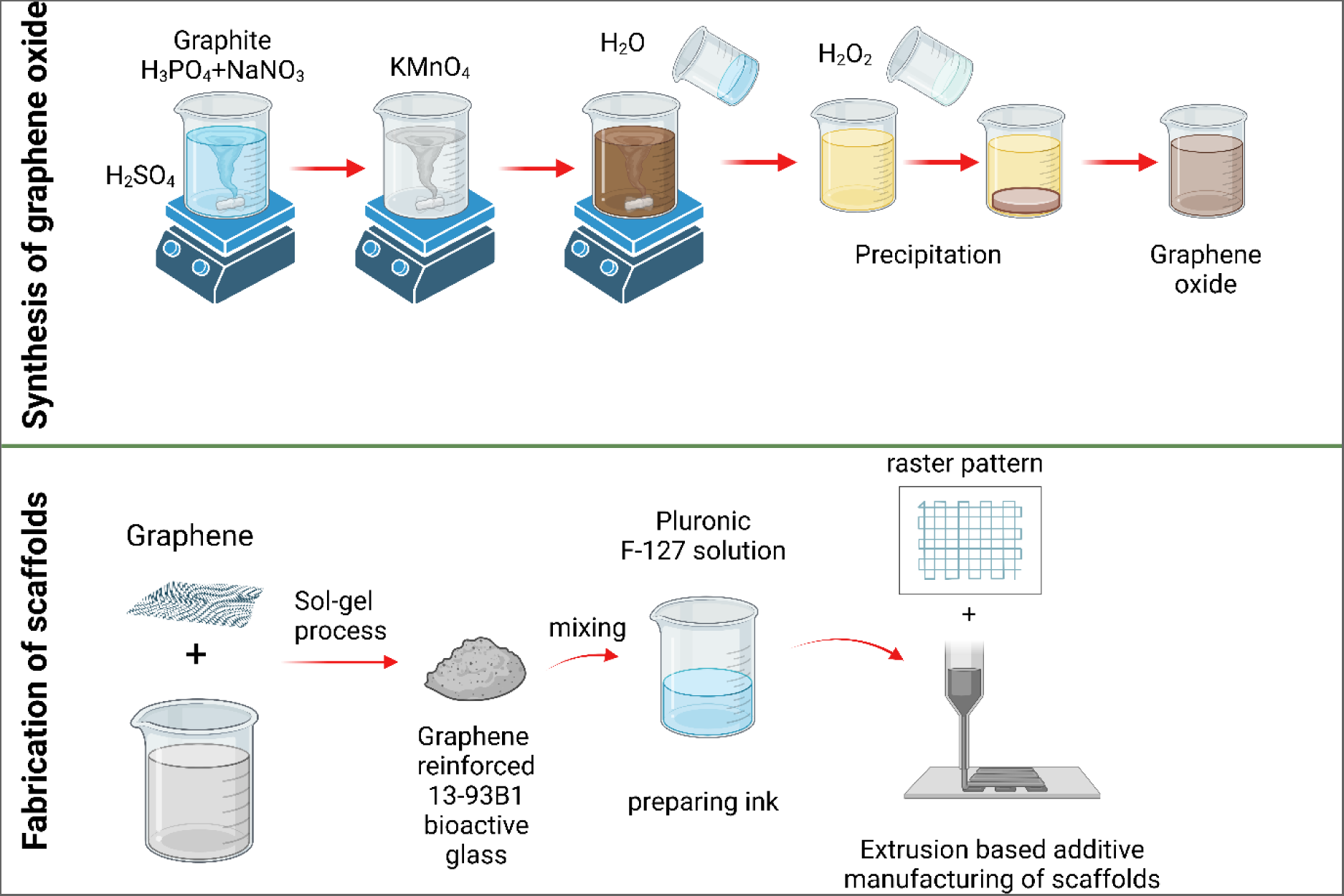
GO synthesis using modified hummer’s method and ink preparation steps for extrusion based additive manufacturing/direct ink writing of the scaffolds.

### 2.4. Direct ink writing of Graphene reinforced bioactive glass scaffolds

To create the ink, an aqueous solution containing 25% Pluronic F-127 (composed of poly (ethylene oxide)-poly (propylene oxide)-poly(ethylene oxide)) was prepared. The 13-93B1 bioactive glass powder was dispersed in the Pluronic F-127 solution using a magnetic stirrer to obtain a paste. It was observed that the paste’s viscosity increased as the bioactive glass input volume percentage increased. As a result, the required extrusion force became extremely high, causing inability to fabricate scaffolds. Insufficient bioactive glass loading, on the other hand, rendered the fabricated struts porous. The fabricated scaffold would therefore exhibit poor mechanical properties. Therefore, the bioactive glass quantity was optimized for direct ink writing and obtaining adequate mechanical properties. The optimum amount was determined to be 38 %(w/v) by volume. In an ice bath, the bioactive glass powder was added in small quantities to the Pluronic F-127 ink. Before the final mixing of the powder, 1 weight percent of octanol-1 was added to the ink to eliminate air pockets. After thoroughly mixing the powder, it was placed in an ice bath for 10 minutes. The GO reinforced bioactive glass ink was prepared in a similar manner.

The ink was subsequently transferred to a 10 mL syringe and used for direct ink writing. by using an in-house developed extrusion-based AM printer. The printing was done at room temperature (23 °C). The raster pattern was provided as G-codes, and Reptier Host was used to control the device. Before commencing the printing of the sample, four additional lines were printed to verify the consistency and flowability of the ink. The diameter of the nozzle used in this study was 410 µm. After printing the sample, it was allowed to dry at room temperature for 24 hours. Depending on the green body, it was then sintered in a muffle furnace (for pure 13-93B1 bioactive glass scaffold) or tubular furnace (for GO reinforced 13-93B1 bioactive glass scaffold).

The green bodies were heated to 400 °C (heating rate 1°C/min), held at that temperature for 1 h, and then sintered at 600 °C for 2 h at a heating rate of 1 °C/min. During the printing process of scaffolds, a layer of nontoxic oleic acid was applied to the printing sheet to prevent crack formation in the green body and to avoid non-uniform sample shrinkage. Additionally, it facilitated the removal of printed scaffolds from the printing platform.

### 2.5. Characterization

#### 2.5.1. Field Emission Scanning Electron Microscopy (FESEM) and Energy Dispersive X-ray Spectroscopy

For evaluation of the microstructure and macrostructure of the fabricated scaffolds in this work. Images were recorded using FESEM (Sigma Carl Zeiss, Oberkochen, Germany). EDS spectra of the scaffolds was also obtained to determine the elemental composition.

#### 2.5.2. Rheological characterization

Constant shear rate flow measurements were performed with a gap of 1 mm between the rheometer platform and the probe geometry at shear rates ranging from 0.01-100 s^-1^ using a rheometer (Anton Paar MCR 302). Dynamic amplitude sweep test ranging from 0.01% to 100% at a constant frequency of 1 Hz was also performed on the ink to evaluate the linear viscoelastic region. The three-interval thixotropy tests, also known as 3ITTs, were carried out in order to evaluate the thixotropic behaviour as well as the phase change that occurred after being subjected to an excessive amount of stress. Shearing the sample at a low shear stress (0.1 Pa) was done for the first and third intervals in order to stabilise the sample and recover the structure, respectively. On the other hand, shearing the sample at a high shear stress was done for the second interval in order to simulate the flow through the nozzle and the resultant structure distortion that occurs during printing.

#### 2.5.3. Micro Computed Tomography (µCT)

In order to investigate the pore size distribution and the interconnectivity of pores, a computed tomography (CT) scan was carried out on the scaffolds with a X-ray CT (X-ray CT mini Procon X-ray GmbH, Germany). The tube voltage was set at 120 kV, and the beam current was set at 100 µA. After acquiring µCT images, they were processed to evaluate pore size pore throat size and porosity of the fabricated scaffold as per the protocol mentioned in Dixit et al [13].

#### 2.5.4. Mechanical property evaluation

The compressive strength of the fabricated scaffolds, 13-93B1 scaffolds and graphene reinforced 13-93B1 scaffolds was evaluated using a universal testing machine. The crosshead speed used during the test was 0.5 mm/min [14]. The compressive strength of the scaffolds was evaluated by dividing the maximum load by cross sectional area of the scaffold.

#### 2.5.5. Perfusion kinetics study of the scaffold

In this study, we have considered a constant inlet velocity having magnitude of 10 µm/s (as this magnitude is considered suitable for osteogenesis) in the normal direction to the boundary, whereas zero pressure boundary condition was specified at the outlet wall [15–19]. No-slip conditions were assumed for all the remaining walls [20,21]. The fluid medium used in this study was blood, as the scaffold is going to be implanted inside the human body. Blood was modelled as both Newtonian and non-Newtonian fluid. Additionally, Dulbecco’s modified eagle medium (DMEM) was also simulated as fluid medium to compare with literature. Permeability (k) of the scaffolds was calculated according to Darcy’s law (with the assumption that the flow is largely one-dimensional) that is given as

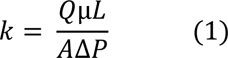

where Q is the fluid flow rate (m^3^/s), µ is the dynamic viscosity of the fluid (Pa.s), L is the length (m), A is the cross-sectional area (m^2^) and ΔP is the pressure difference between inlet and outlet face of the scaffold (Pa). Grid independence tests were also conducted to get optimal grid (please refer to Figure S1). The difference between the grid having number of elements 577232 and 414572 is less than 2%. Therefore, we have taken the grid having 414572 number of elements for the CFD simulation of all cases. For permeability calculations, the properties of blood were considered with a dynamic viscosity of 0.00345 Pa.s and density of 1.05 g/cm^3^. Further, the pressure drop for permeability was calculated for two cases namely by considering blood as a Newtonian fluid and non-Newtonian fluid, respectively. Overall permeability of the scaffold was considered to be the addition of 𝑘_𝑥_, 𝑘_𝑦_, and 𝑘_𝑧_. 𝑘_𝑥_ and 𝑘_𝑦_ were permeability values reported parallel to the printing plane while 𝑘_𝑧_ was permeability measured in the z-direction i.e., perpendicular to the printing plane. For flow simulations, the blood has been assumed to be a non-Newtonian fluid and its dynamic viscosity has been incorporated into the model using Carreau-Yasuda model that is expressed as follows:

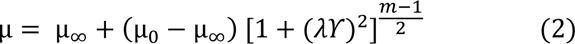

where µ_0_ indicates low shear viscosity with a magnitude of 0.056 Pa.s, µ_∞_ represents high shear viscosity having a magnitude of 0.00345 Pa.s, λ represents the time constant (λ = 3.3313 s), m being the power law index having a value of 0.3568 [22].

#### 2.5.6. *In vitro* biocompatibility

The C2C12 mouse myoblast cells were used as model mammalian cells for evaluating the biocompatibility of additively manufactured GO reinforced 13-93B1 bioactive glass scaffolds. The C2C12 cells were cultured using a cell growth media comprising Dulbecco’s modified Eagle’s medium (Lonza, Basel, Switzerland) supplemented with 10% heat-inactivated fetal bovine serum (Thermo Fisher Scientific, Waltham, Massachusetts) and 1% penicillin– streptomycin (Sigma-Aldrich). The cells were incubated in at 37°C and 5% CO_2_ atmosphere till further use.

Next, the scaffolds were sterilized under UV radiation for 1 hour and placed in a 24-well plate with 1 scaffold per well. The C2C12 cells were treated with trypsin-EDTA solution (Sigma-Aldrich) to detach from the tissue culture flask and collected as a pellet by centrifuging at 1500 rpm for 3 minutes. The pellet was further resuspended in fresh cell culture medium. Each scaffold containing well was loaded with approximately 5000 cells and 2mL culture media and maintained at 37°C with 5% CO_2_. The viability of the cells was evaluated after 1, 2 and 3 days of culture. First, the samples were transferred to a Petri dish and gently rinsed twice with PBS to remove the cell culture media. Next, the samples were immersed in live/dead assay (Biotium, Hayward, CA, USA) solution for 30 minutes at room temperature. Next the live/dead assay was aspirated gently followed by rinsing with PBS and immersion in PBS for further imaging. The assayed samples were imaged using a fluorescent microscope (Revolve, ECHO, CA, USA). Two different fluorescent channels corresponding to Fluorescein isothiocyanate (FITC) and Texas red were used to image Calcein-AM stained live cells and EthD-III stained dead cells respectively. The cell viability was evaluated by counting the number of cells in the FITC (𝑛_𝐹𝐼𝑇𝐶_) channel, Texas red (𝑛_𝑇𝑒𝑥𝑎𝑠 𝑟𝑒𝑑_) channel images and overlapping cells (𝑛_𝑜𝑣𝑒𝑟𝑙𝑎𝑝_) in both images respectively using particle counter plugin in ImageJ (NIH, USA) and equation Eq. 1.

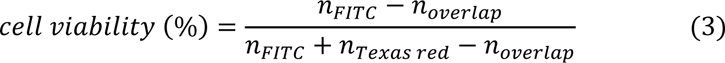

The morphology of the cells adhered on the scaffolds was examined by scanning electron microscopy (SEM). After 3 days of culturing with C2C12 cells, the scaffolds were washed three times with PBS. Next the scaffold samples were fixed by immersing in 4% v/v glutaraldehyde for three hours and finally rinsed three times with PBS for ten minutes to remove residual glutaraldehyde. Next, the scaffolds were dehydrated with varying graded concentrations of ethanol (10-100% v/v) for 15 minutes and vacuum-dried overnight. The scaffolds were mounted on stubs using a carbon tape and imaged using an SEM (Phenom Pro X, Thermo Scientific, MA, USA).

Cell proliferation was evaluated using the XTT (2,3-Bis-(2-methoxy-4-nitro-5-sulfophenyl)-2H-tetrazolium-5-carboxanilide) assay (Biotium Inc., Fremont, CA) according to the manufacturer’s instructions. Two representative sets were prepared for comparison by culturing C2C12 cells in cell growth media without the scaffold (control) and with the scaffold (13-93B1 Bioactive glass scaffold with GO reinforcement). In a 24-well plate, an initial population of 5000 C2C12 cells was seeded on each of the two sample categories. The samples were maintained at 37°C with 5% CO_2_ and 1 mL cell growth media in an incubator. To allow for initial cell adhesion and attain confluency, all samples were incubated for 48 hours before the spent cell growth medium was replaced with fresh cell growth medium. Following the addition of 1 mL of fresh cell culture media to each well, 100 µL of activated XTT solution was added. The samples were then incubated for 4 hours to permit the formation of dark red formazan salt crystals, which are indicative of metabolic activity within the cells. Next, 200 µL of the solution was transferred from each well of the 24-well plate to a 96-well plate. The absorbance of each solution was measured at a wavelength of 450 nm using a UV–vis microplate reader (Bio-Rad, Philadelphia, PA).

## 3. Results and Discussion

### 3.1. Particle size distribution, FESEM, EDS and Rheological characterization

The size of the bioactive glass particles plays a crucial role in determining the compressive strength of the scaffolds. As the particle size decreases, the compressive strength of the scaffolds increases. Conversely, larger particle sizes can lead to higher ink viscosity, making it unsuitable for printing. To address this, the bioactive glass powder was subjected to ball milling using zirconia balls and subsequently sieved. Figure 3A shows the variation of volume % vs particle size distribution of 13-93B1 bioactive glass. In this study, the D_50_ value was measured to be 4 μm. Also, the D_90_ value was determined to be 19 μm. Furthermore, the volume-weighted mean particle size was calculated as 7 μm.

**Figure 3.**
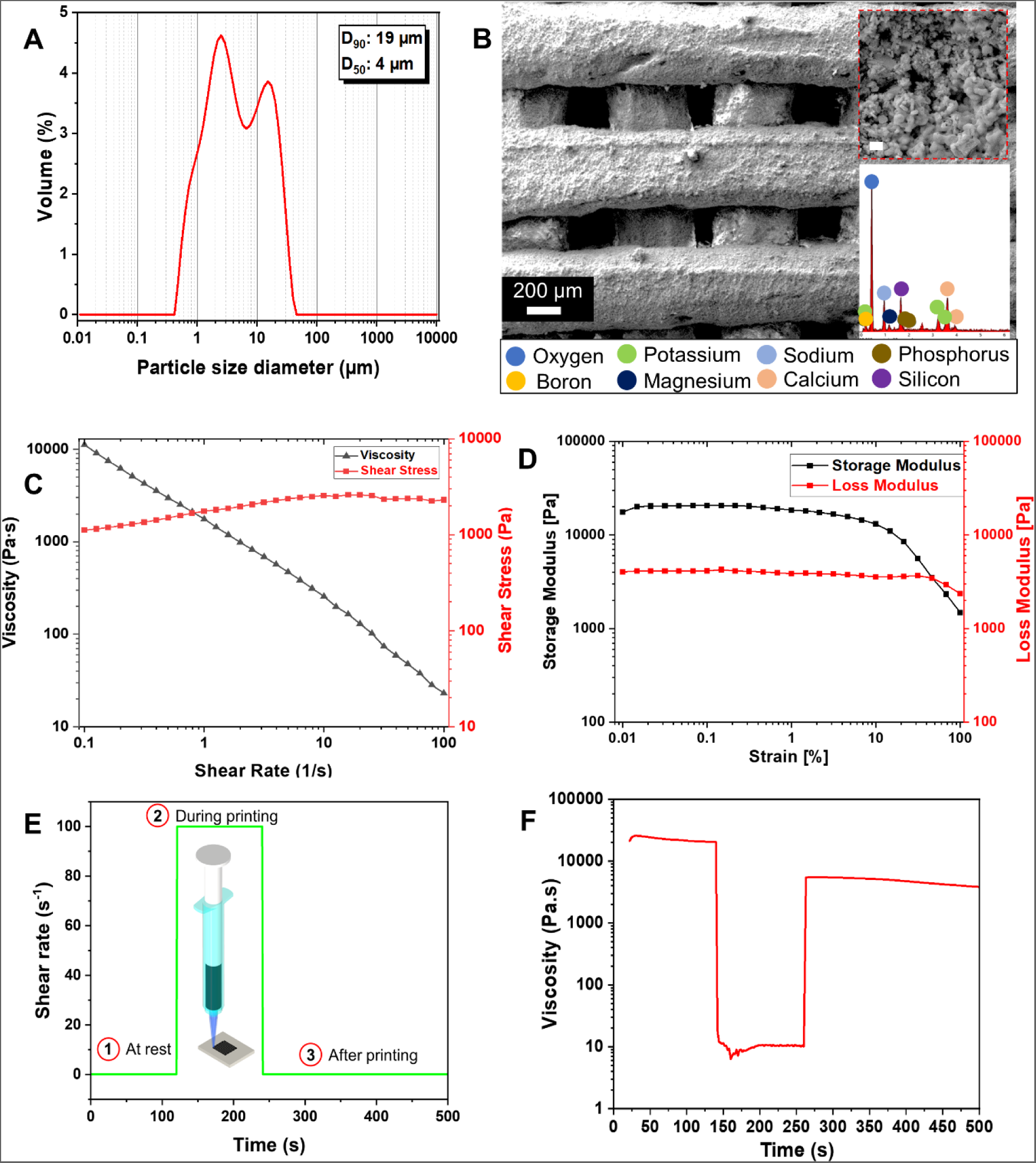
(a) Particle size distribution, (b) FESEM and EDS of 13-93B1 bioactive glass scaffold, (c) variation of shear stress and viscosity with shear rate, (d) amplitude sweep test result of ink (e) testing conditions of 3 interval thixotropy test and (f) 3ITT result showing shear thinning behaviour and structural recovery behaviour of the ink.

Figure 3B shows the FESEM image of the fabricated scaffold showing the wood pile structure. The inset of the FESEM image shows the porous strut of the fabricated scaffold. It also shows the elemental composition of the scaffold as obtained from the EDS spectra.

Figure 3C shows variation of viscosity and shear stress with shear rate. The shear thinning behaviour of the GO-reinforced bioactive glass ink was modelled using the Herschel Bulkley model [23]. The following equation describes the Herschel Bulkley model:

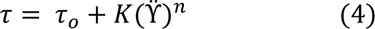

where 𝜏 denotes shear stress, 𝜏_𝑜_expresses the yield shear stress, K symbolizes the viscosity parameter of the ink, ϔ correspond to shear rate and n represents the flow index, which is also known as the shear thinning exponent. If the value of n is below 1, the studied ink exhibits shear-thinning behavior, meaning its viscosity decreases with increasing shear. Figure 3C illustrates the shear stress vs. shear rate data, fitted into the Herschel-Bulkley model, yielding n and K values of 0.18±0.02 and 1725±279, respectively. This affirms the suitability of GO-reinforced 13-93B1 bioactive glass ink for additive manufacturing of scaffold. The molecular mechanisms for shear-thinning and shape retention vary among ink types. For instance, polymers experience shear-induced disentanglement of long polymer chains, while for dispersions and pastes, shear-thinning results from the breakdown of solid particle interactions. The ink’s shape stability post-extrusion is attributed to its high viscoelasticity at rest, reestablishing contacts between suspended particles. Yield stress, the stress required to initiate flow, indicates the ink’s shape retention; a higher yield stress suggests better shape retention due to increased cross links or entanglements within the ink, enhancing scaffold’s final stiffness and strut formation [24]. Furthermore, dynamic sweep test was conducted on the GO-reinforced 13-93B1 bioactive glass ink to evaluate flow behaviour (Figure 3D). Initially the storage modulus shows higher value as compared to loss modulus at smaller strain %. The storage modulus (G’) shows elastic behaviour i.e. solid or gel like behaviour (G’ > G”) of the ink while loss modulus (G”) refers to fluid like behaviour (G’ < G”). With increasing strain % there is a cross over of G” over G’ indicating flow initiation of the ink.

Moreover, 3ITT was conducted to analyse the ink’s performance throughout different stages of additive manufacturing [25]. Ensuring self-supporting 3D structures with good shape fidelity relies on the transition kinetics resembling fluid flow and exhibiting solid-state behaviour. Rapid recovery of the printed ink is essential to prevent continuous flow post-extrusion, restoring elastic behaviour. This test also evaluates the ink’s response to various shear rates, ranging from low shear rates during rest after loading to high shear rates during nozzle flow. The ink’s ability to recover after printing, experiencing very low shear rates, is crucial for maintaining structural integrity and supporting subsequent layers. Figure 3F illustrates the ink’s viscosity variations during these simulated steps, indicating its suitability for additive manufacturing, demonstrating load-bearing capacity, shape retention, and support for successive layers.

### 3.2. Raman spectra

Figure 4 shows the Raman spectra of graphene oxide (GO), 13-93B1 bioactive glass and GO reinforced 13-93B1 bioactive glass. GO’s Raman spectrum displays two main characteristic peaks at 1356 and 1599 cm^-1^, which correspond to the D and G bands, respectively. The D band is the result of sp^3^ defects, whereas the G band is the result of in-plane vibrations of sp^2^ carbon atoms and a doubly degenerated phonon mode [49]. Due to the clustering of GO sheets, these peaks become less intense and broader. Further, Raman spectrum of 13-93B1 bioactive glass shows Si-O-Si peak at 1050 cm^-1^ referring to stretching vibration. In case of GO reinforced 13-93B1 bioactive glass scaffold, the typical peaks associated with both GO and 13-93B1 bioactive glass were observed. Si-O-NBO (Non bridging oxygen) peak is observed at 960 cm^-1^ indicating the disruption of the silica network [26].

**Figure 4.**
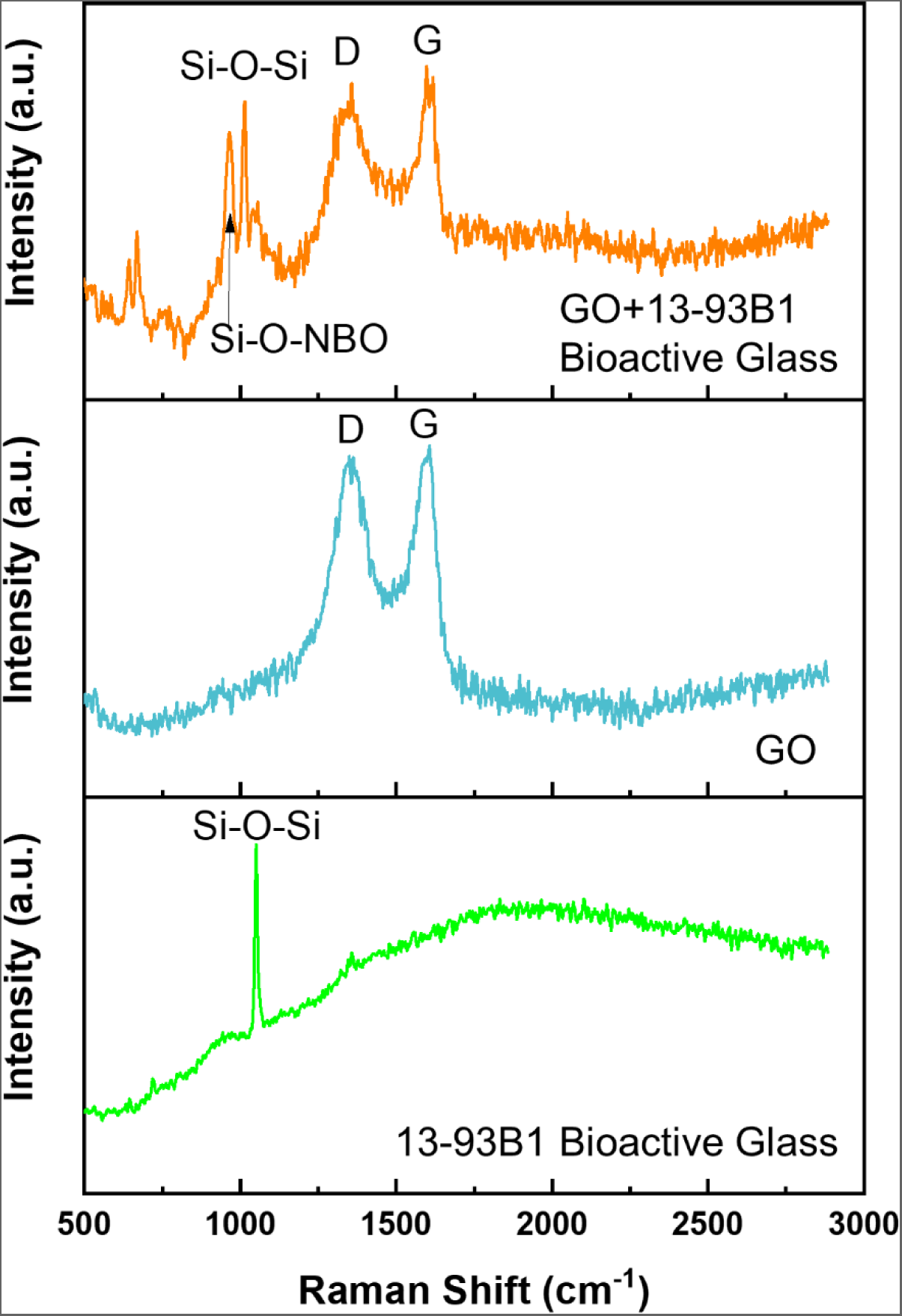
Raman spectra of (a) 13-93B1 Bioactive Glass, (b) Graphene oxide, and (c) GO-reinforced 13-93B1 bioactive glass scaffold.

### 3.3. Morphological characterization using micro-computed tomography

Figure 5A shows the bioactive glass matrix region, while pore region is presented in Figure 5B acquired from the µCT images. The 3D reconstruction of the matrix region and pore region is presented Figure 5D and 5E, respectively. Pore size and pore throat size distribution of the fabricated scaffold obtained as per the procedure mentioned in section 2.5.5. Pore size and pore throat size distribution of the fabricated scaffold are presented in Figure 5C and Figure 5F, respectively. The fabricated GO reinforced 13-93B1 scaffolds were found to have a mean pore size of 460 µm and a throat diameter of 157 µm, both of which are larger than the 100 µm limit typically cited in the literature [27,28]. Therefore, the scaffolds’ porous structure is adequate for nutrient delivery and metabolic waste disposal. Also, most bone cell sizes (20-30 µm) may reach the whole scaffold without any path restriction, as shown by the interconnectivity analysis, which demonstrates 99 % accessibility with a structural element diameter of size 80 µm for GO reinforced 13-93B1 scaffolds. Figure 5G shows the pore throat network model of the scaffold. The network shows the grid like connections of the pores that is similar to the SEM images of the fabricated scaffold. Figure 5H is a visualization of the AM-facilitated through-and-almost-straight routes present in the constructed scaffold. These results suggest that manufactured scaffolds have the necessary morphological features for use in bone tissue engineering.

**Figure 5.**
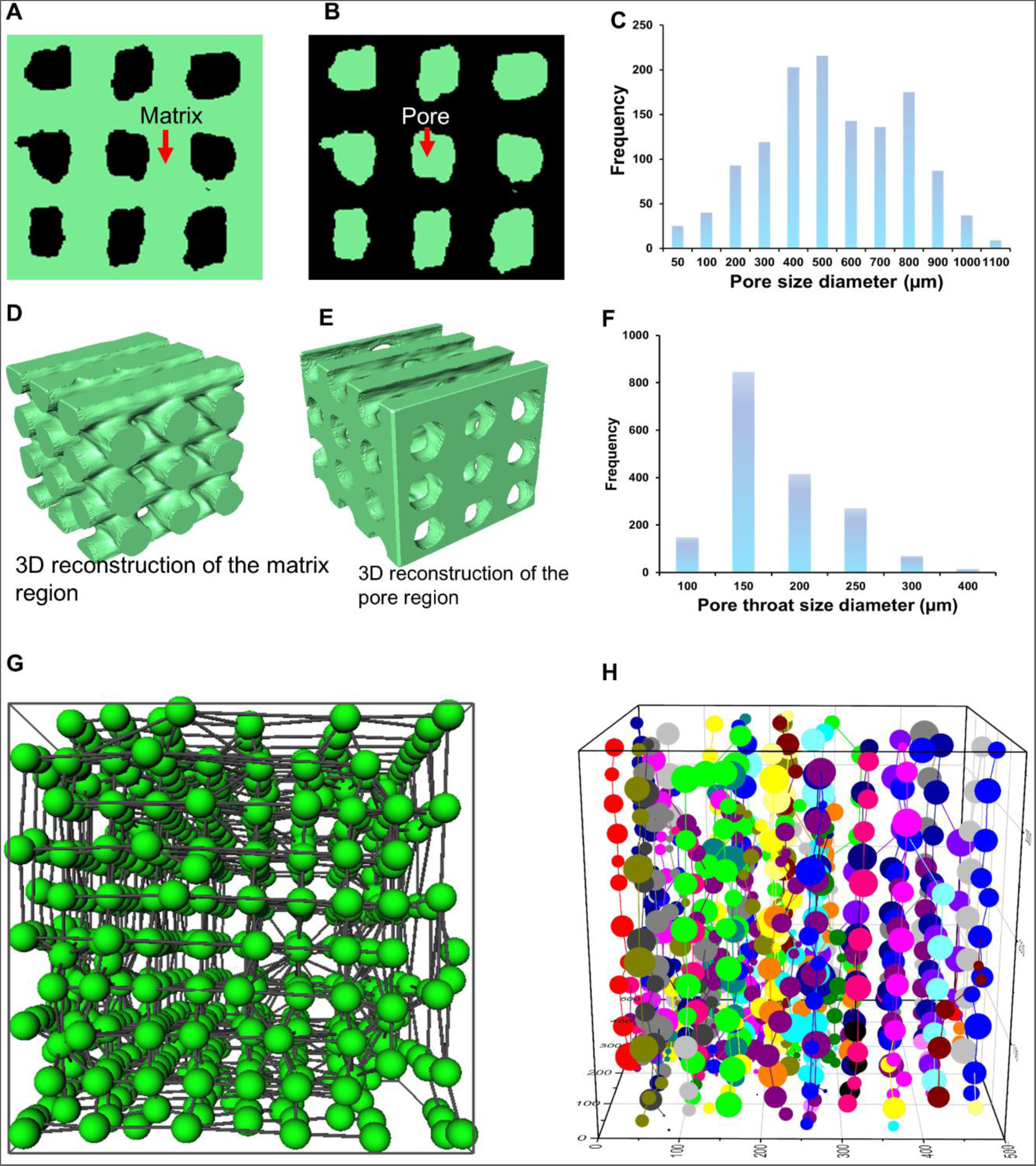
(a) µCT image of a subvolume showing matrix region and pore region in (b), (c) pore size distribution,(d) and (e) 3D reconstruction of matrix and pore region, (f) pore throat size distribution, (g) top view of pore network model and (h) through path visualization in the GO-reinforced 13-93B1 bioactive glass scaffold.

Further, the interconnectivity is measured as the ratio of connected pore volume to total pore volume for a given structural element diameter [29]. Figure S2 depicts the variation of interconnectivity with varying structural element diameter values. It is evident that 99 % of the scaffold structure is accessible by a structural element (SE) having 120-µm diameter. Therefore, it can be concluded that the majority of bone cell sizes (20–30 µm) have unrestricted access to the entire scaffold. However, as the size of the SE increases, it consequently decreases the scaffold’s accessibility (pore interconnectivity). With an increase in the diameter of the SE, the interconnections (throats) are blocked, resulting in a reduction in pore interconnectivity. With each iteration, an increasing percentage of interconnections between pores were disconnected, reducing the proportion of the total connected pore volume. With an adequate SE diameter of 332 µm for molecular transport, the scaffold’s accessibility continued to be 50 %. Thus far, it is possible to conclude that the fabricated scaffold has a well-connected pore network.

### 3.4. Mechanical characteristics of the fabricated scaffolds

Figure 6A shows the compressive stress vs compressive strain curve of the pure 13-93B1 bioactive glass and GO reinforced 13-93B1 bioactive glass scaffolds. The compressive strength of the fabricated scaffolds evaluated using the methods outlined in section 2.2 is presented in

**Figure 6.**
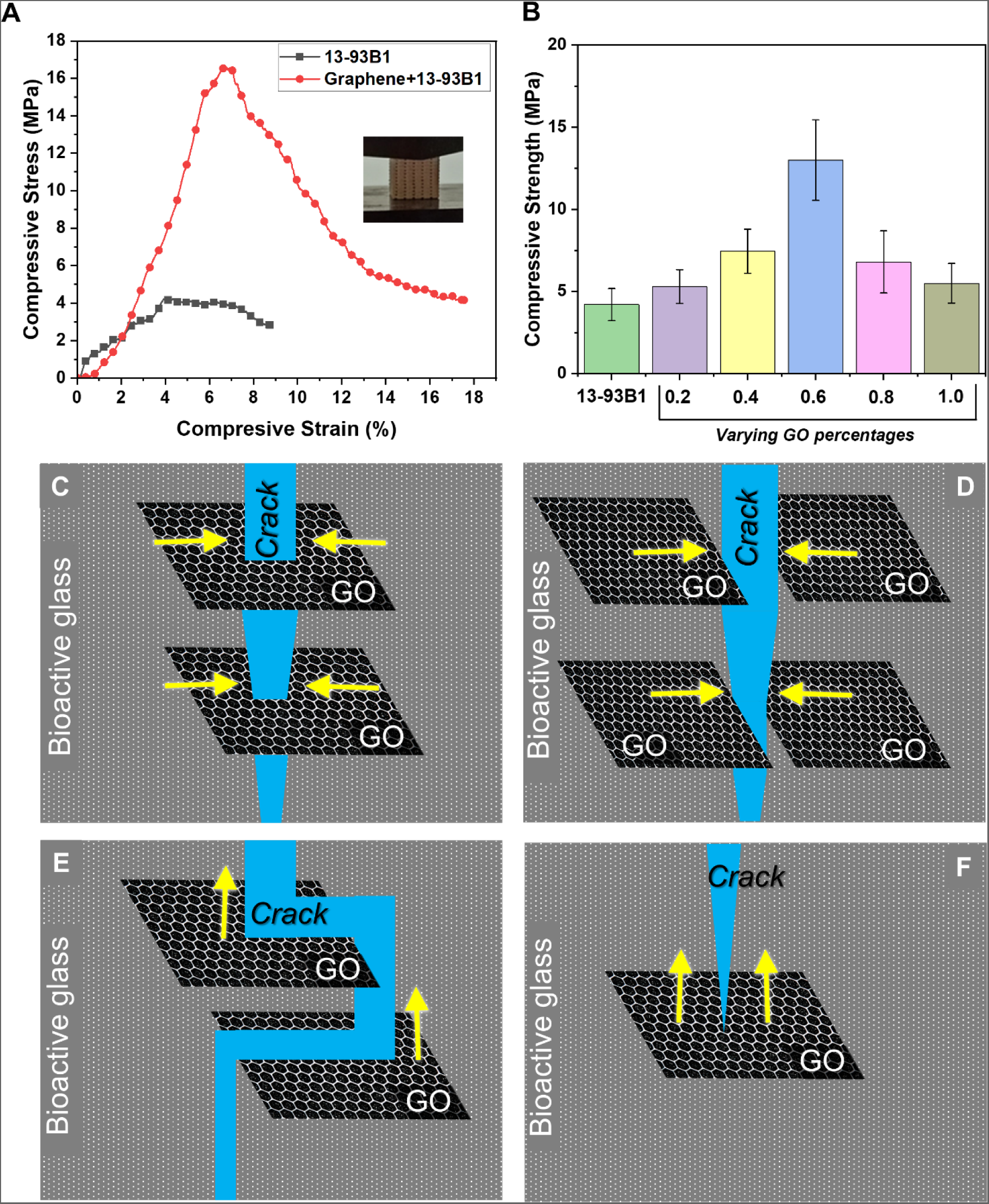
(a) Compressive stress vs. compressive strain of 0.6 weight percentage GO reinforced 13-93B1 bioactive glass scaffold and (b) comparison of compressive strength of the scaffold having varying percentages of GO with pure 13-93B1 bioactive glass scaffold, Schematic showing (a) pull-out, (b) crack bridging, (c) crack deflection, and (d) crack tip shielding mechanism by GO dispersed in 13-93B1 bioactive glass.

Figure 6B. It can be seen that with increasing strain, the compressive stress of the scaffold increases. However, the stress-strain curve of the scaffolds exhibited an initial elastic response before reaching the maximum load bearing capacity of the scaffold. After reaching the maximum load capacity, crack appeared on the unsupported struts close to the supported sturts of the scaffold. Further, more cracks appeared throughout the struts of the scaffold, the compressive strength continued to decrease until the densification of the fractured scaffold, which is a usual characteristic of cellular scaffolds. However, the addition of GO to 13-93B1 bioactive glass significantly enhanced the scaffold’s resistance to damage (i.e., its toughness), resulting in a larger area under the compressive stress vs strain curves (please refer to Figure 6 A).

Further, Figure 6B show that with increasing concentration of GO, the compressive strength of the scaffolds increases, however, after a certain amount of enhancement the reinforcement in compressive strength decreases. Further, it was found that 0.6 % (w/w) of the GO resulted in maximum compressive strength enhancement (∼12.7 MPa). The compressive strength values of the scaffold having 0.6 % (w/w) of the GO is comparable to the compressive strength of the trabecular bone [30]. These enhancements can be explained using the mechanisms such as crack bridging, crack deflection, uniform load transfer (please refer to Figure 6C-F [31,32]. The special characteristics of GO and its interactions with the bioactive glass matrix are responsible for these mechanisms. The aforementioned mechanisms are discussed in more detail as follows:

GO’s remarkable mechanical properties stem from its unique two-dimensional structure, which consists of a single layer of carbon atoms arranged in a hexagonal lattice. The uniform dispersion of GO across the scaffold, creates a network of reinforcements [33]. Further, GO has ability to bear load and transfer stress due to its strong covalent bonds. Consequently, these capabilities of GO contribute to the scaffold’s enhanced structural integrity and compressive strength as a whole. Additionally, the inclusion of GO promotes efficient load transfer from the scaffold. The efficient transfer of stress from the bioactive glass matrix to the GO sheets is made possible by the strong interaction between GO and the bioactive glass matrix [34]. This mechanism redistributes the applied load throughout the scaffold, increasing its compressive strength, and prevents localised stress concentrations [35].

In the scaffold, GO also functions as a barrier, preventing cracks from propagating further inside the bioactive glass matrix. Crack growth and propagation are slowed in GO sheets because of their two-dimensional structure, high aspect ratio, and exceptional mechanical properties. Compressive forces cause cracks to propagate in the bioactive glass matrix. Moreover, GO acts as a bridge, reinforcing the weak points, and increasing the scaffold’s fracture toughness [31,36].

Strong interfacial bonding and interaction between GO and the bioactive glass matrix are essential to the scaffold’s integrity. The bioactive glass particles can exhibit strong covalent and non-covalent interactions with the GO sheets due to the presence of functional groups on the surface of GO sheets, such as hydroxyl and carboxyl groups. Overall, improved load transfer and stress distribution are outcome of this interfacial bonding, leading to enhanced compressive strength of the scaffold [37].

### 3.5. Perfusion kinetics of the fabricated scaffold

The permeability of a scaffold is pivotal in bone tissue regeneration and influences nutrient, oxygen delivery to cells and waste removal from the scaffold. Generally, in a scaffold, two mechanisms facilitate the movement of metabolites to cells and the removal of waste products: diffusion and, in the context of *in vivo* application, transportation facilitated by capillary networks developed within the scaffold through angiogenesis. Although angiogenesis poses a limiting factor in *in vivo* situations, noteworthy angiogenesis does not occur during the initial days post-implantation and is entirely absent in *in vitro* environments [38]. Further, it has been reported that a higher permeability for a scaffold may result in enhanced bone regeneration as higher permeability may promote more osteogenic related protein adsorption due to availability of more attachment sites [39,40].

Hui et al. [41] investigated the cancellous bone graft’s permeability. They concluded that a minimum threshold permeability of 3×10^-11^ m^2^ is required for angiogenesis and bone mineralization inside the implant. The authors also concluded that a high permeability value in a scaffold may result in its enhanced integration with the native tissue. From the CFD study of the fluid flow through the scaffold in this work (please refer to Figure 7, 8 and 9), our estimated permeability of the GO reinforced 13-93B1 bioactive glass scaffold (2.08×10^-9^ m^2^) is found comparable to the values reported in previous studies for synthetic bone grafts (see Table 2). We noted that our fabricated scaffolds have superior transport properties since the permeability value is comparable to most of the previous studies presented in Table 2. The scaffolds in this table that have permeability comparable to the present work, also have comparatively higher porosity. The permeability of the GO reinforced 13-93B1 bioactive glass scaffold indicates the potential for uninhibited vascularization and better nutrient transport to the interior regions of the scaffold. Therefore, the computed permeability values prove the scaffold’s potential for tissue regeneration.

**Figure 7.**
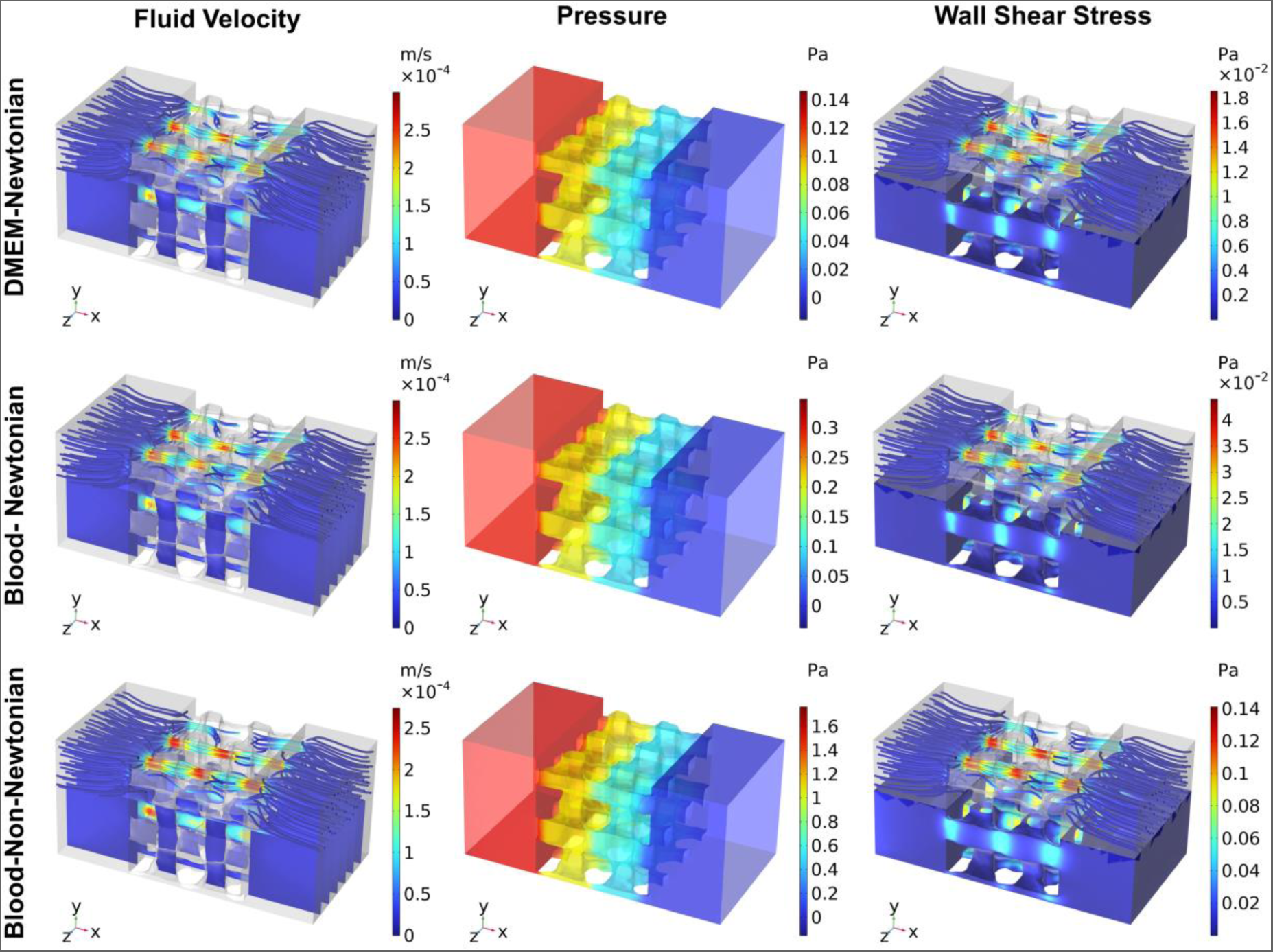
(a)Velocity (slice and streamline distribution), (b) pressure, and (c) wall shear stress distribution (surface shows the wall shear stress distribution along with the velocity streamlines) in the scaffold with a fluid velocity of 10 µm/s in X-direction.

**Figure 8.**
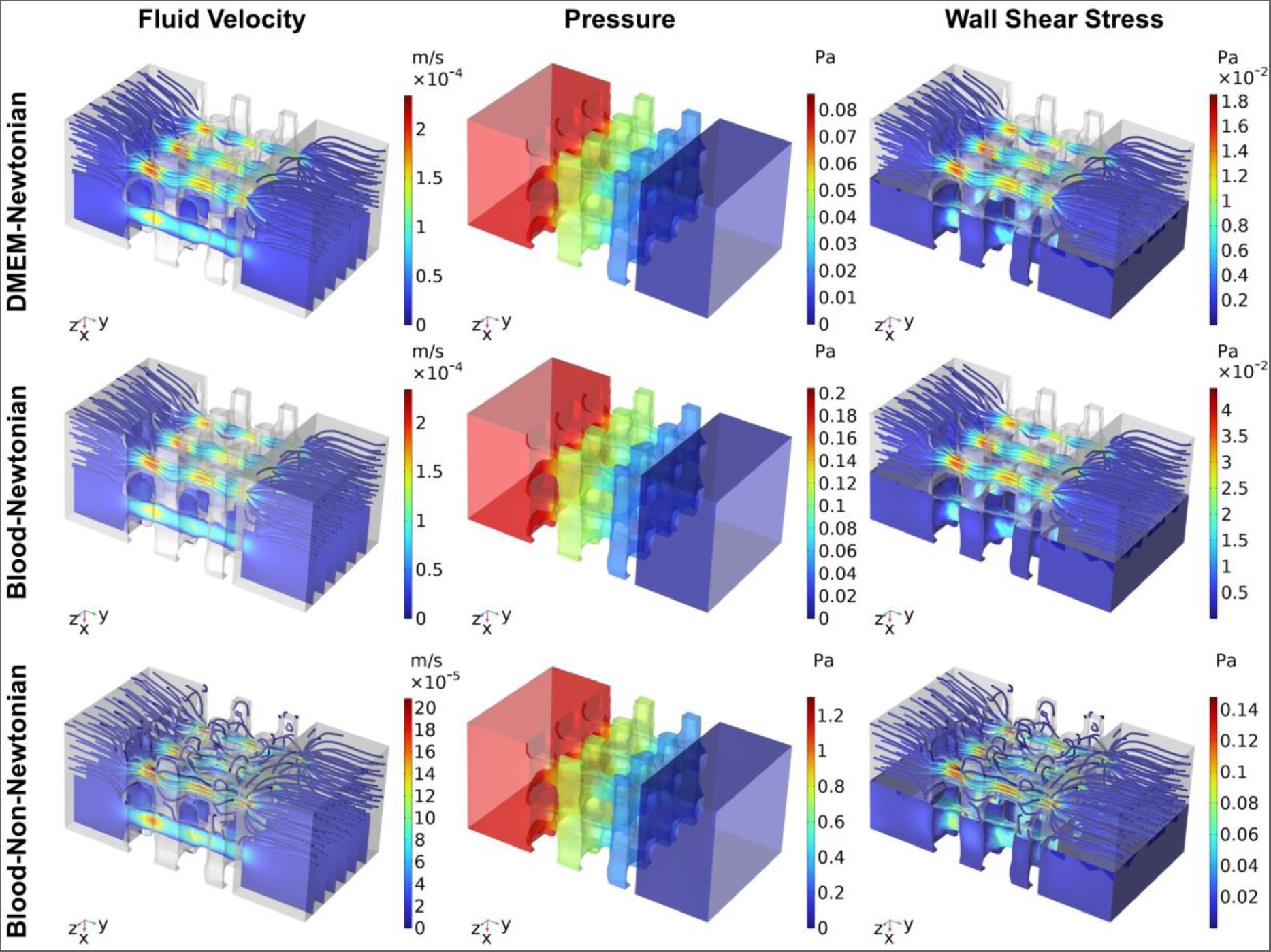
(a)Velocity (slice and streamline distribution), (b) pressure, and (c) wall shear stress distribution (surface shows the wall shear stress distribution along with the velocity streamlines) in the scaffold with a fluid velocity of 10 µm/s in Y-direction.

**Figure 9.**
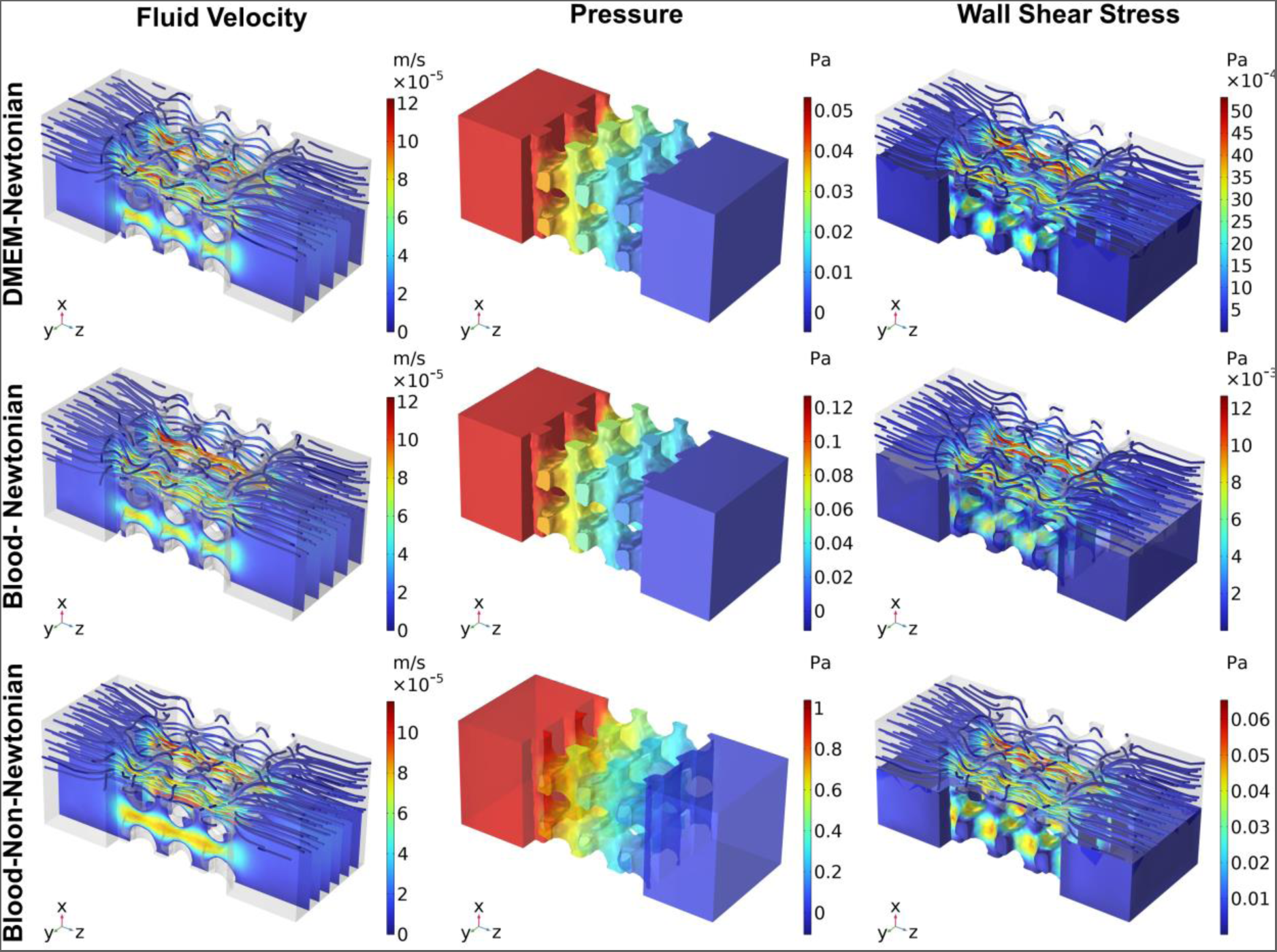
(a)Velocity (slice and streamline distribution), (b) pressure, and (c) wall shear stress distribution (surface shows the wall shear stress distribution along with the velocity streamlines) in the scaffold with a fluid velocity of 10 µm/s in Z-direction.

**Table 2.** Comparison of average wall shear stress and permeability of various scaffolds.

| Material of the scaffold | Porosity (%) | Mean pore size ($\mu\text{m}$) | Inlet velocity ($\mu\text{m/s}$) | Average Wall shear stress (mPa) | Permeability ($\text{m}^2$) | Reference |
| --- | --- | --- | --- | --- | --- | --- |
| Polyurethane | 77 | 100 | 53 | 3.28-3.94 | $3.1 \times 10^{-12}$ | Cioffi et al. 2006 [42] |
| Calcium phosphate | ~32 | 300-400 | 10 | 37 (max) | - | Sandino et al. 2008 [15] |
| Glass | 30 | - | 10 | 46.55(max) | - |  |
| Titanium | 77 | 280 | 34 | 1.41 | - | Maes et al. 2012 [43] |
| Hydroxyapatite | 73 | 270 | 34 | 1.09 | - |  |
| Bio-Oss | 60 | 320 | 10 | 1.35 | $3 \times 10^{-10}$ | Zhang et al. 2019 [44] |
| Cerabone | 69 | 300 | 10 | 1.44 | $6 \times 10^{-10}$ | |
| Maxresorb | 67 | 140 | 10 | 1.98 | $0.31 \times 10^{-10}$ | |
| Sr doped $\text{Ca}_2\text{ZnSi}_2\text{O}_7$ (Architecture A) | 49.7 | 390 | - | - | $7.9 \times 10^{-10}$ | Entezari et al. 2019 [45] |
| Cancellous Bone | 50-90 | - | | | $2.68 \times 10^{-11}$-$2 \times 10^{-9}$ | Nauman et al. 1999 [46] |
| 13-93B1-GO* | 55 | 460 | 10 | 1.3 (z-direction)<br>2.6 (y-direction)<br>3.8 (x-direction) | $1.07 \times 10^{-9}$ (z-direction)<br>$1 \times 10^{-9}$ (printing plane)<br>$2.08 \times 10^{-9}$ (Overall)* | This work |

\*Blood considered as Newtonian fluid

Fluid velocity distribution in the porous region of the scaffold is visualized using a combination of slice distribution and streamlines. The flow pattern of the fluid in the scaffold showed an increase in the fluid velocity when pore cross section area decreased, while with expanding area a drop in fluid velocity was observed. The fluid velocity has direct implications for nutrients and oxygen transport. Proper distribution of fluid flow helps in homogeneous distribution of nutrients, ensuring uniform growth of cell.

Wall shear stress contours of the scaffold corresponding to different flow directions viz., x, y and z are shown in Figures 7, 8 and 9. The growth of cells and their behaviour in the scaffolds is affected by wall shear stress magnitude [47]. When a scaffold is implanted into the body, it comes into contact with blood that flows over the surface of the scaffold. This blood flow on the surface of the scaffold results in force that is called wall shear stress. Various studies have reported that magnitude of the wall shear stress may result in different effects on cells as depicted in Table 3. In this work we have simulated three cases, DMEM as Newtonian fluid, blood as Newtonian and blood as non-Newtonian fluid using Carreau Yasuda model. The obtained average wall shear stress values were compared with the critical wall shear stress values reported in the literature. Overall, the obtained average wall shear stress values were found to have a stimulating effect on the cells if cultured under the simulated conditions.

**Table 3.** Magnitude of critical wall shear stress for cells in a scaffold having culture media as fluid during perfusion (Adapted from Maes et al. [43])

| **WSS (mPa)** | **Effect on cells** | **References** |
| --- | --- | --- |
| 1.5-13.5 | Stimulation | Raimondi et al [48] |
| 0.05 | Proliferation | Cartmell et al. [49] |
| 1 | Differentiation |  |
| 57 | Apoptosis |  |
| 4.6 | Stimulation | Raimondi et al. [50] |
| 14-25 | Differentiation |  |
| 56 | Cell wash out |  |
| 1.1-3.8<sup>\$</sup> | Stimulation | This **work** |
| 10.3-13.2<sup>#</sup> |  |  |

**\$DMEM and Blood as Newtonian Fluid, #Blood as non-Newtonian fluid**

It is evident from the previous studies (Table 3) that a very high magnitude of the wall shear stress may result in cell death or detachment/wash out while lower magnitudes are preferable for cells leading to their stimulation, proliferation and differentiation. Further, it has been reported in the literature that the cell tend to align themselves along the fluid flow direction in the regions of high wall shear stress also resulting in enhanced proliferation [51].

The wall shear stress values obtained in this work with the computational fluid dynamics simulations performed using DMEM and Blood as fluid (Newtonian and Non-Newtonian). When modelled as Newtonian fluid the wall shear stress values vary directly in proportion to the viscosity. For instance, blood having higher viscosity (0.00345 mPa.s) as compared to DMEM (0.00145 mPa.s) showed increased magnitude of wall shear stress (1.5 to 3.8 mPa along X-direction). Additionally, in case of pressure distribution, blood modelled as Newtonian fluid has more resistance than the culture media leading to higher magnitude of pressure at the inlet part of the flow. Further, the pressure distribution in each flow direction showed higher magnitude in case of non-Newtonian fluid than the Newtonian fluids indicating that the non-Newtonian fluid has more resistance to flow.

### 3.6. *In vitro* results: cell proliferation and cell viability

After characterizing the physical and chemical properties of the GO-reinforced 13-93B1 bioactive glass scaffolds, we evaluated their biocompatibility *in vitro* to assess their potential as implantable scaffolds. Once implanted, these scaffolds are expected to interact with surrounding bone as well as muscle tissues. In this study we focused on the bone-muscle interface and, therefore, assessed the compatiblity of developed GO-reinforced 13-93B1 bioactive glass scaffolds with C2C12 mouse myoblast cells. Live dead assay reavealed high biocompatibility of the GO-reinforced 13-93B1 bioactive glass scaffolds with C2C12 cells when cultured on the surface of the scaffolds as shown in Figure 10A. Majority of the scaffold surfaces were covered with the cells with a nearly homogeneous distribution. When cultured for 3 days, the scaffolds maintained their biocompatibility beyond 95% (Figure 10B) indicating that release of various ions from the scaffolds did not hinder the cell growth. A closer observation also showed elongated and branched morphology of the cells characteristic of adherent cell types like C2C12. This implied the presence of cell-adhesive capabilities in the developed GO-reinforced 13-93B1 bioactive glass inks as well as their homogenous distribution within the bioprinted scaffolds. Cell adhesiveness is an important property in bone tissue engineering as it governs the integration of the implanted scaffold with native bone and muscle tissues towards bone regeneration [52,53].

**Figure 10.**
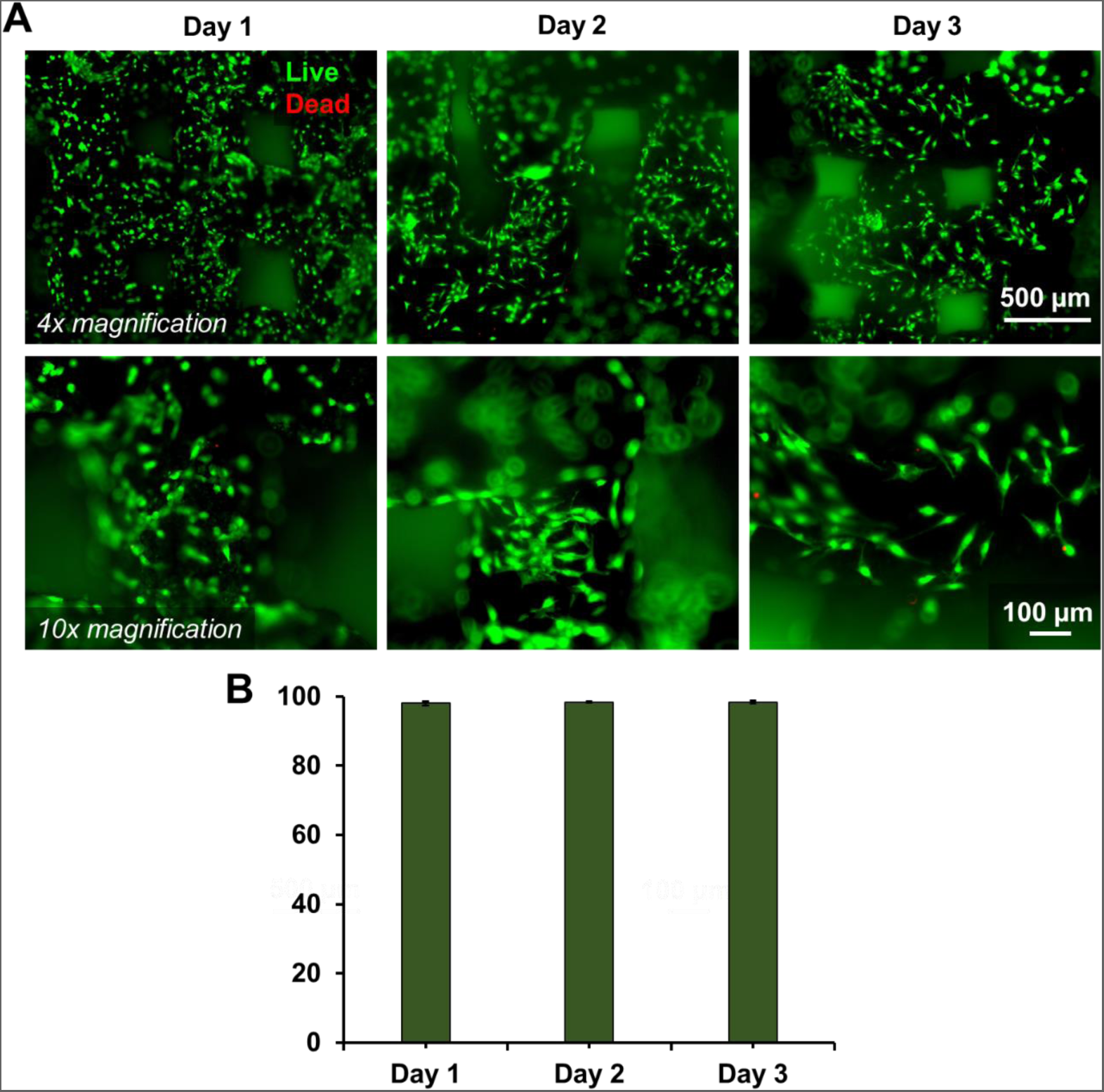
Live/dead assay results of C2C12 cells after day 1, day 2, and day 3 cell culture on GO-reinforced 13-93B1 bioactive glass scaffolds.

The cells attached on the scaffold surface were further examined unsed SEM after 3 days of culture. The struts in the cell-laden scaffolds exhibited a porous surface with presence of small crystal-like structures (Figure 11) similar to the strut of the GO reinforced 13-93B1 bioaactive glass scaffolds which demonstrated a compact porous surface (Figure 3B). Formation of small crystal-like structures indicates formation of hydroxyapatite (HAP) as shown in previous studies [54]. Closer examination of the cell-scaffold interface in the SEM images (Figure 11 insets) reveals HAP to be possible binding sites for the C2C12 cells. However, the cell-adhesion mechanism for C2C12 on GO-reinforced 13-93B1 bioactive glass scaffolds will be explored in detail in future studies.

**Figure 11.**
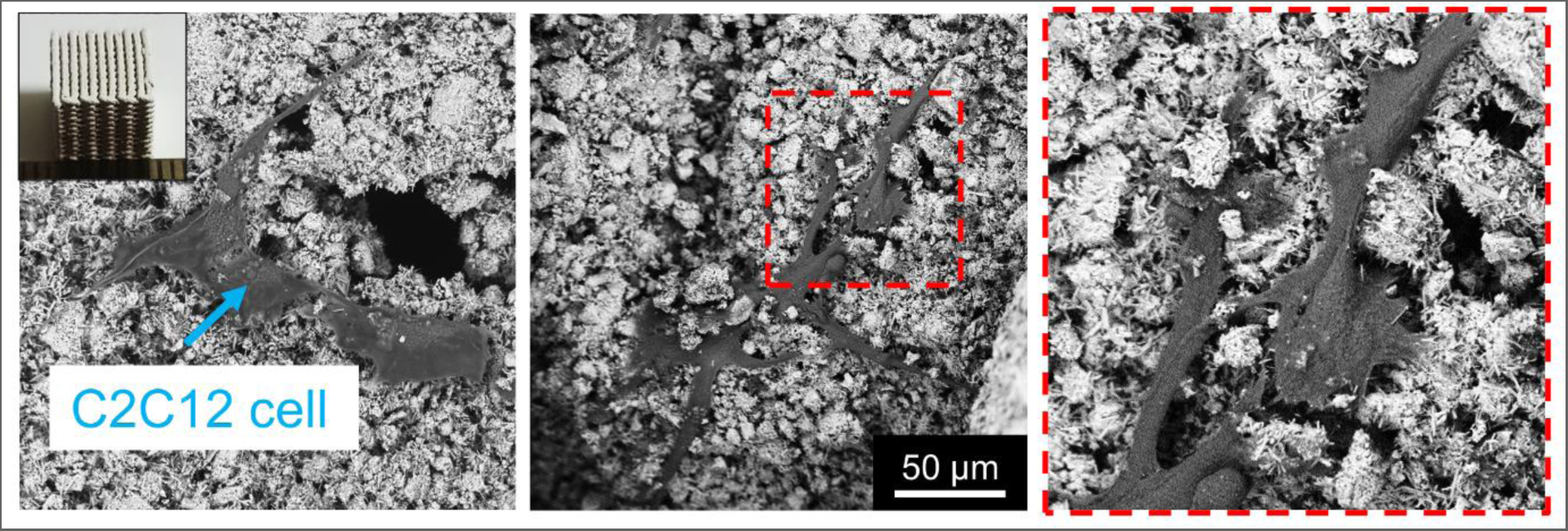
SEM images of GO-reinforced 13-93B1 bioactive glass scaffold showing cell adhesion of C2C12 cells after three days of cell culture (Inset showing image of the additively manufactured scaffold).

Beyond cell viability, we also assessed the influence of the developed BG-Gr scaffolds on the growth of the C2C12 cells. The scaffolds were introduced to the cells cultured on a 12-well plate with varied contact durations. Metabolic activity of the cells was considered as representative of the proliferation of the cells and was quantified using XTT assay(Figure 12). The cells were also observed to grow in the proximity of the scaffolds and often atttempting to form cell-cell networks bridging the structs of the scaffolds (Figure 12A). The morphology of the cells in absence and presence of the scaffold remained unchanged (Figure 12B). The XTT assay demonstrated a slight increase in the overall metabolic activity upto 18 hours of contact with the scaffolds (Figure 12C). When observed beyond 18 hours and up to 48 hours, the overall metabolic activity remained consistent with slight decline. This observation can be attributed to the wells getting overpopulated with the cells and slowing down the overall metbolism.

**Figure 12.**
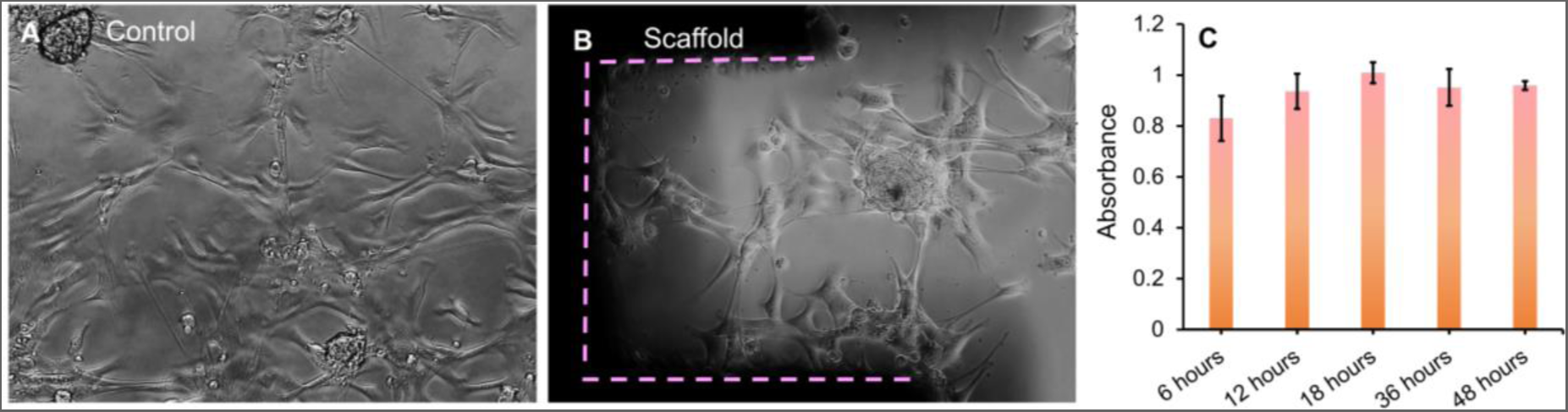
(a) GO-reinforced 13-93B1 bioactive glass scaffold with C2C12 cells after 48 hours of cell culture, (b) C2C12 cells in tissue culture plate (control), and (c) Cell proliferation evaluated using XTT assay.

## 4. Conclusions

In this study, we present the development of a GO-reinforced 13-93B1 bioactive glass scaffold using additive manufacturing for bone tissue engineering applications. The scaffold fabrication process involved the incorporation of GO nanoplatelets into a bioactive glass powder, followed by the extrusion-based 3D printing of the composite ink by using an in-house developed AM machine. The GO reinforcement resulted in enhanced mechanical properties of the scaffold allowing it to withstand physiological loads. The fabricated scaffolds exhibited a well-defined interconnected porous structure, mimicking the natural architecture of bone tissue. Moreover, the perfusion kinetics of the scaffold obtained using CFD simulations exhibited its suitability of permeability in the range of trabecular bone. Additionally, the WSS levels obtained using simulations were found to be suitable for stimulating cell growth. Obviously, it warrants further experimentation in near future to validate these results. Furthermore*, in vitro* evaluations demonstrated the bioactivity of the GO-reinforced 13-93B1 bioactive glass scaffold, cell viability and proliferation studies revealed the scaffold’s biocompatibility, with enhanced cell adhesion and growth of C2C12 cells.

Overall, the developed GO-reinforced 13-93B1 bioactive glass scaffold fabricated using additive manufacturing holds great promise for bone tissue engineering applications. Its tailored architecture, mechanical strength, and bioactivity make it a suitable candidate for promoting bone regeneration and facilitating cell-material interactions in the perspective of bone tissue engineering.

## Acknowledgement

The authors would like to thank Science and Engineering Research Board, Department of Science and Technology, India for partially supporting this study through grant number SB/FTP/ETA-464/2012. Kartikeya would like to thank Alberta Innovates for Alberta Innovates Postdoctoral Fellowship.

