## Supplementary Information for "Direct Ink Writing of Graphene Oxide Reinforced 13-93B1 Bioactive Glass Scaffolds for Bone Tissue Engineering Applications"

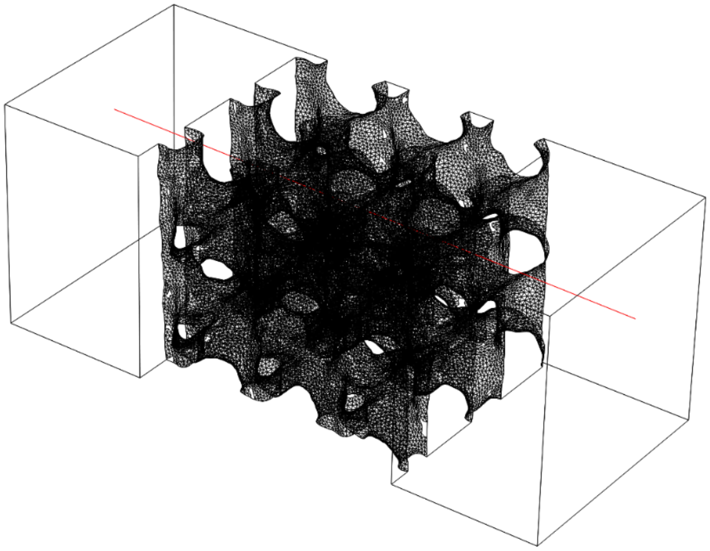

(a)

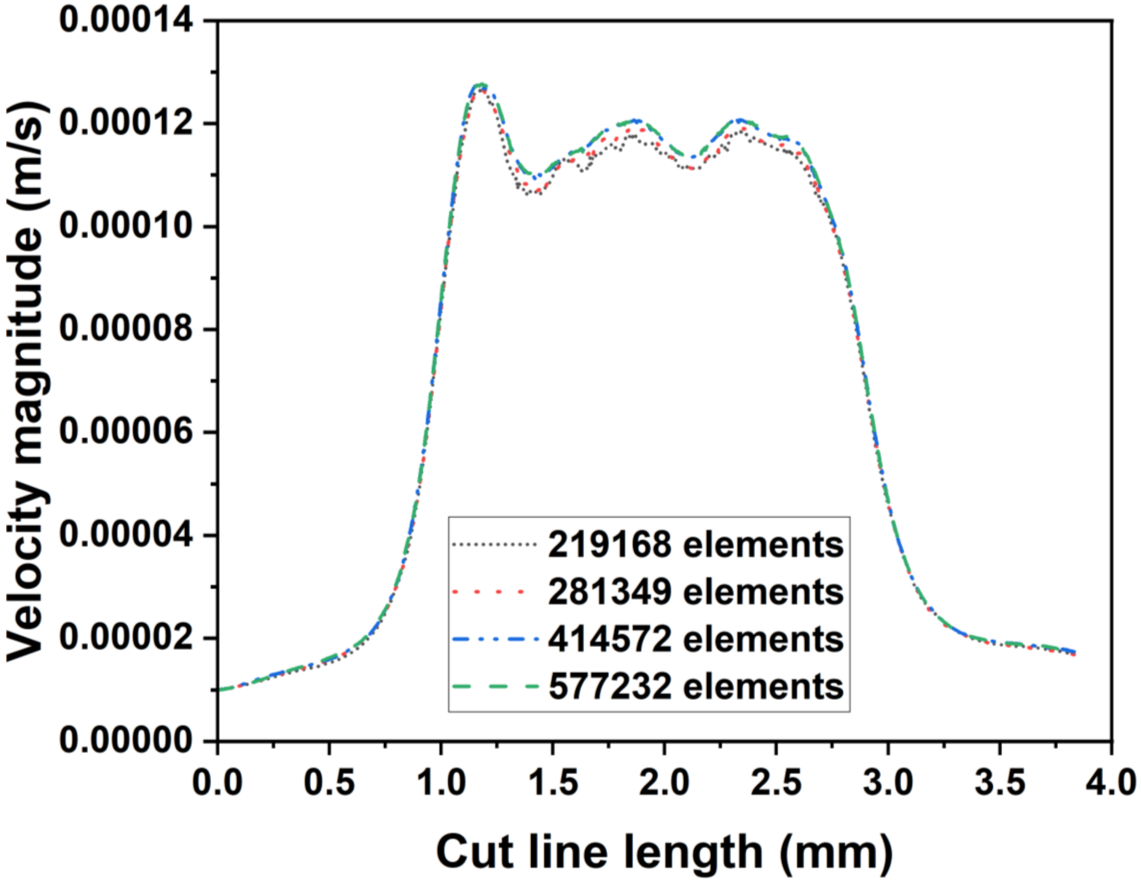


(b)

Figure S1. (a) location of the cut line in the fluid domain, and (b)Variation of fluid velocity with number of elements along the cut line showing mesh sensitivity.

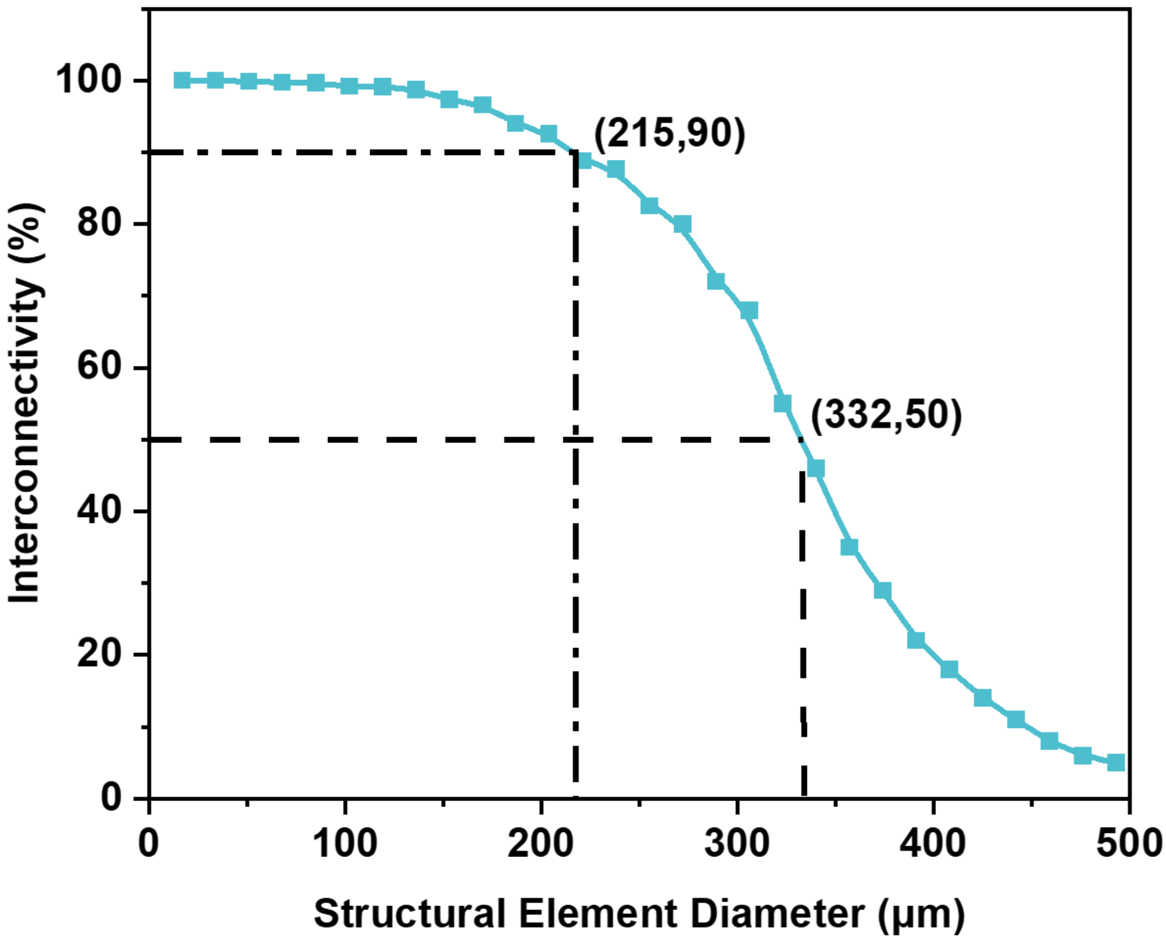


Figure S2. Interconnectivity variation with structural element diameter for GO-reinforced 13-93B1 bioactive glass scaffold.
